# Dietary iron overload enhances susceptibility to *Yersinia enterocolitica* infection

**DOI:** 10.64898/2026.08.24.746810

**Authors:** McKenzie Van der Veer, Shreya Das, Nicole Vienneau, Dapeng Zhang, Wei Sun

## Abstract

Hemochromatosis and hemosiderosis are iron overload disorders that cause immune dysfunction and increase susceptibility to bacterial infections. There have been numerous case studies reporting septic-like outcomes for hemochromatosis patients infected with enteric Yersiniae; however, research regarding hemosiderosis and *Yersinia* infection is limited. Here, we have established a mouse model of hemosiderosis by feeding C57BL/6 mice a high-iron diet. These mice exhibit several indicators of iron overload that are seen clinically, including elevated serum iron levels and iron deposition in various tissues. Characterization of the iron overload mouse model shows that a high-iron diet induces local inflammation in the small intestine and systemic inflammation in a time-dependent manner. Oral infection with *Yersinia enterocolitica* causes complete mortality in the iron-overloaded mice, while wild-type mice all survive and effectively clear the infection. Lastly, we have observed that iron chelation therapies such as Deferoxamine and Deferisarox are detrimental to iron-overloaded mice during *Yersinia* infection. This work provides a model to further study iron overload disorders and *Yersinia* infection.

**Importance:** Iron overload disorders can lead to severe organ dysfunction, immune system dysregulation, and increased risk of infection. Specifically, there have been numerous case studies reporting septic-like outcomes for hemochromatosis patients infected with enteric Yersiniae. Here we have established a dietary iron overload mouse model which can be utilized to characterize the impacts of high iron levels on immunity and *Yersinia* infections. Advancements in this field will be beneficial for improving treatment strategies for infections in individuals with hemochromatosis and hemosiderosis.

## Introduction

Hemochromatosis and hemosiderosis are disorders of iron metabolism characterized by pathological iron overload (IO), leading to iron deposition in various tissues and resulting in complications such as liver disease, cardiomyopathy, cancer, immune dysfunction, and increased susceptibility to infections. The term hemochromatosis refers to IO of genetic origin (1), caused by mutations affecting hepcidin production (2). These mutations lead to hepcidin deficiency, increased intestinal iron absorption, elevated plasma iron levels, and iron deposition in parenchymal cells (3–5). The term hemosiderosis refers to other types of tissue IO, which can be developed through excessive iron supplementation, hemolysis, or frequent blood transfusions (3, 6). In contrast to hemochromatosis, hemosiderosis typically exhibits increased hepcidin, leading to iron accumulation within phagocytic cells (3, 5).

It is known that IO predisposes individuals to infections caused by a broad spectrum of pathogens, including bacteria, fungi, and viruses (7, 8). Numerous case studies have reported septic-like outcomes in hemochromatosis patients infected with the enteric Yersiniae, including *Y. enterocolitica* (Ye) and *Y. pseudotuberculosis* (Yptb) (9, 10). Both Gram-negative pathogens can infect humans through ingestion of contaminated food or water (11). In the small intestine, Ye and Yptb utilize microfold cells to cross the intestinal epithelial barrier (IEB) and reach the Peyer’s Patches (11). The bacteria migrate to the mesenteric lymph nodes and establish a local infection, manifesting as gastroenteritis (11). Ye infection is common in Western countries and typically mild and self-limiting in healthy individuals, but can cause 20-50% mortality in IO conditions (12). Similar to what is observed in humans, the Ye Ruokola/71 (serotype O:9) strain, a clinical isolate (13), was avirulent in wild-type (WT) C57BL/6 mice but lethal in hepcidin-1 knockout (HKO) mice (14).

Oral iron supplementation is the most common treatment for iron deficiency anemia (IDA) and non-anemic iron deficiency (ID), which are among the most common pathologies worldwide (15, 16). Iron repletion using oral iron supplements require high doses for prolonged periods of time (16). Only a fraction of this iron is absorbed, while the excess accumulates in the gastrointestinal (GI) tract, leading to intestinal inflammation and increased susceptibility to infection (16). Further, the use of nutritional supplements has increased due to growing public interest in healthy lifestyle (17). Iron supplementation has become very common, and iron supplement users have been shown to exceed the recommended dietary allowance (17). The most commonly used oral iron supplement is ferrous sulfate (16). In a study involving human subjects, a single dose of ferrous sulfate was shown to be sufficient to induce oxidative damage in healthy individuals (17, 18). As the use of oral iron supplementation rises, it is important to further investigate its impacts on the GI system.

Human studies have utilized iron-fortified diets to investigate how high iron levels can impact the intestine and infection. It has been shown that high dietary iron favors the growth of pathogenic bacterial species, which is accompanied by intestinal inflammation (19). Also, oral iron supplementation has been shown to significantly increase the risk of hospitalization and mortality associated with various infections (20–22). It has been reported that pathological patterns and complications associated with nutritional IO in humans are similarly observed in dietary IO mice (23), and the dietary IO model appears to be analogous to human hemosiderosis (24). The dietary IO mouse model, in which mice are fed a high iron diet (HID), has been used to examine iron-induced pathology (25) and *Salmonella* infection (26). However, there has been little work done to evaluate the effects of dietary IO in the context of Ye infection.

In this study, we have investigated how dietary IO affects iron homeostasis, systemic and intestinal immune response, and survival upon Ye infection. We found that feeding mice a 2% carbonyl iron diet induces hemosiderosis, triggers systemic and intestinal inflammation, and results in pronounced susceptibility to Ye infection. The dietary IO mouse model established here will facilitate investigation of the underlying mechanisms by which high iron levels predispose hosts to Ye infection, a clinically significant condition due to its prevalence in the United States and worldwide, thereby supporting the development of effective therapeutic strategies to combat bacterial infections in affected individuals.

## Materials and Methods

### Mice and animal ethics statement

All animal studies were performed per the NIH “Guide for the Care and Use of Laboratory Animals” and approved by the Institutional Animal Care and Use Committee at Albany Medical College (IACUC protocol# 25-12001). Animal experiments were performed using WT C57BL/6 (B6) mice obtained from the Jackson Laboratory and bred in our animal facility. Six- to eight-week-old male and female B6 mice were fed a 2% Carbonyl Iron 10% kcal Fat Diet (93G) (TD.230570, Tekland) for two weeks or four weeks to generate a HID mouse model termed 2wk Fe and 4wk Fe mice, respectively. The composition of the HID is outlined in Table S1.

### Iron and hepcidin assays

The serum iron concentration and serum transferrin saturation were determined using the Total Iron-Binding Capacity and Serum Iron Assay Kit (Cayman Chemical, Catalog No. 702230). The serum ferritin concentration was determined using a Mouse Ferritin ELISA Kit (Invitrogen, Catalog No. EEL095). All assays were performed according to the manufacturer’s instructions.

### Iron staining

Tissue iron deposition was evaluated by Perls’ Prussian Blue staining (25, 27) using the Iron Stain kit (Sigma, Catalog No. HT20). Briefly, tissue sections of liver, small intestine, heart, kidney, and spleen were deparaffinized and rehydrated in distilled water, incubated in iron stain (1:1 potassium ferrocyanide solution: hydrochloric acid solution, 1 hr, 25°C), rinsed in three washes of distilled water, and incubated in pararosaniline solution (1:50 pararosaniline solution: distilled water, 5 min, 25°C). The slides were rinsed in three washes of distilled water before rapidly dehydrating in alcohol and xylene and mounting.

### Detection of ferrous iron in spleen and liver mononuclear phagocytes (MNPs)

B6, 2wk Fe, and 4wk Fe liver and spleen samples were processed, stained for extracellular markers, and incubated in BioTracker FerroOrange Live Cell Dye (Sigma-Aldrich, Catalog No. SCT210) prior to detection by flow cytometry using a BD FACSymphony A3 (BD Biosciences). The data was analyzed using FlowJo. Briefly, cell strainer pestles were used to break up the spleen or liver and pass the cells through 70 µm strainers. The strainers were continuously flushed with 0.5% BSA in 1x phosphate buffered saline (1× PBS) (FACS buffer). Splenic or liver cells were centrifuged (400 x g, 6 min., 4 °C), the supernatant was discarded, and the cell pellet was disrupted by vortexing. The cells were incubated in TheraPEAK Ammonium-Chloride-Potassium (ACK) Lysing Buffer (Lonza Bioscience, Catalog No. BP10-548E) for red blood cell lysis (3 mL, 3 min., 25°C), neutralized with FACS buffer (9 mL), and centrifuged (400 × g, 6 min., 4 °C). After the supernatant was discarded, splenic cells were resuspended in FACS buffer (3mL) and passed through another 70 µm strainer, while liver cells were resuspended in FACS buffer (4mL) and passed through a 40 µm strainer.

Intracellular ferrous iron (Fe^2+^) was detected by flow cytometry. 3×10^6^ live cells/sample were centrifuged and resuspended in anti-mouse CD16/32 (1:500, 50 µL, 3-5 min., 4°C) prior to incubation with antibodies to extracellular markers (50 µL, 40-60 min., 4°C). The samples were neutralized with FACS buffer (900 µL), centrifuged (400 × g, 6 min., 4 °C), resuspended in FerroOrange (200 µL, 1µM), and incubated (30 min., 37°C) prior to detection by flow cytometry.

### Complete blood count (CBC)

Whole blood was collected from B6, 2wk Fe, and 4wk Fe mice. Complete blood count analysis was performed using a Heska Element HT5 (Heska, Colorado).

### Cytokine analysis Bio-Plex Pro assay

Serum samples and intestinal homogenates were assessed for various cytokines using the Bio-Plex Pro Mouse Cytokine Grp 1 Panel 23-Plex (Bio-Rad, Catalog No. M60009RDPD). To make intestinal homogenates, sections of the small intestine (∼50 mg) were homogenized using a Bertin Precellys 24 tissue homogenizer, centrifuged (8,000 x g, 10 min., 4°C), and the supernatant was collected and stored in -80°C. The assay was performed according to the manufacturer’s instructions, the plate was read using a Bioplex-200 reader (Bio-Rad, USA), and the data were analyzed using the Bioplex Data Pro Software v (Bio-Rad, USA).

### Hematoxylin and eosin (H&E) staining

Tissue sections of the small intestine were stained using the H&E Stain Kit (Abcam, Catalog No. ab245880). Briefly, sections were deparaffinized and rehydrated in distilled water, incubated in hematoxylin (5 min., 25°C), rinsed in two changes of distilled water, dipped in bluing reagent (15 sec., 25°C), rinsed again, and dipped in absolute alcohol prior to incubating in eosin (10 min., 25°C). Lastly, slides were rinsed in absolute alcohol, rapidly dehydrated in alcohol and xylene, and mounted. Images were obtained using a NanoZoomer S60 (Hamamatsu Photonics, Japan) and analyzed by NDP. View2 software.

### Claudin-3 Immunohistochemistry (IHC)

Tissue sections of the small intestine were deparaffinized and rehydrated in distilled water. Slides were incubated in citrate buffer for antigen retrieval (20 min., ∼100 °C), then transferred to ice cold distilled water (5 min., 4°C). Then the tissues were permeabilized in 0.4% Tween20 (20 min., 25°C), and washed in 0.05% Tween20 (3x, 2 min./wash, 25°C) prior to blocking. 5% Bovine Serum Albumin (BSA) in wash buffer was used as blocking buffer and was added to each tissue (100 µL, 2 hrs., 4°C). After blocking, the slides were washed as previously described, and primary antibody, anti-mouse Claudin-3 (raised in Rabbit), was added to the tissues and incubated overnight (1:300, 100 µL, 4°C). The next day the slides were washed, and secondary antibody (anti-Rabbit A647), was added (1:400, 100 µL, 1 hr., 4°C). The slides were washed again and mounted. Images were obtained using a Zeiss LSM 880 confocal microscope.

### Claudin-3 Western Blot

Samples were prepared by homogenizing B6, 2wk Fe, and 4wk Fe small intestine samples in RIPA Buffer containing a protease inhibitor (1:100) as previously described. The total protein concentration per sample was quantified using the Pierce BCA Protein Assay Kit (ThermoFisher Scientific, Catalog No. 23225), and 2.5 µg/µL stocks were made to normalize the protein concentration for each sample prior to loading and running SDS-PAGE. The protein bands were then transferred onto nitrocellulose membranes and blocked with a 5% skim milk blocking buffer (overnight, 4°C). After blocking, the membrane was washed three times with 1% Tris-buffered saline and tween 20 (TBST) followed by one wash with 1 × PBS. The membrane was then cut between 25-35 kDa. The membrane containing protein bands <25-35 kDa was incubated with primary antibody Anti-Claudin 3 (raised in rabbit), while the membrane containing protein bands >25-35 kDa was incubated with anti-β-actin antibody (raised in mouse) (2 hrs., 37°C). The membranes were then washed as previously described and HRP-conjugated anti-rabbit and HRP-conjugated anti-mouse secondary antibodies were added to the anti-claudin 3 and anti-β-actin blots respectively (1 hr., 37°C). The nitrocellulose blots were washed, developed using ECL™ Detection Reagents (Sigma, USA), and the protein bands were visualized and imaged using the ChemiDoc XRS+ Imaging system (Bio-Rad, USA). The ratio between Claudin 3 and β-actin was quantified using ImageJ software.

### Fluorescein isothiocyanate (FITC)-Dextran (FD4) Assay

The FD4 assay was used to measure intestinal permeability. Food was removed from B6, 2wk Fe, and 4wk Fe mice for 4-6 hours prior to treatment with FD4 (Sigma Aldrich, Catalog No. FD4). The mice were left access to water to prevent dehydration. After the fasting period, mice were treated with 150 µL of FD4 (80mg/mL) by oral gavage and remained on a water-only diet. Four hours after FD4 treatment, blood was collected from the mice via submandibular bleeding into tubes containing Ethylenediaminetetraacetic acid (EDTA) (15 µL, 0.5 M). Plasma was collected by centrifuging the whole blood (1,500 × g, 12 min., 4 °C) and collecting the top layer. The FD4 fluorescence in the plasma was measured using a BioTek Synergy HT Microplate Reader (BioTek, Winooski, VT) at an excitation of 485/20 nm and emission of 528/20 nm.

### Intestine processing and flow cytometry

The small intestine was isolated and processed for flow cytometry as described previously (28), with minor modifications. The Peyer’s patches, fat, and fecal matter were removed, and the intestine was cut open longitudinally for better exposure to the digestion buffer. Intestine samples were chopped into small pieces and incubated in a digestion buffer containing 4-(2-hydroxyethyl)-1-piperazineethanesulfonic acid (HEPES) (20 mM), liberase (166 µg/mL), and DNase (100 µg/mL), in RPMI 1640 medium (3 mL/sample, 30 min., 37 °C). The samples were neutralized with FACS buffer, centrifuged (400 x g, 6 min., 4 °C), resuspended in FACS buffer (1 mL), and passed through a 70 µm strainer. Samples were centrifuged, resuspended in FACS buffer (1 mL), passed through a 40 µm strainer, centrifuged, and resuspended in FACS buffer (1 mL).

For analysis by flow cytometry, 3×10^6^ live cells/sample were centrifuged and resuspended in anti-mouse CD16/32 (1:500, 50 µL, 3-5 min., 4°C) prior to incubation with antibodies (50 µL, 40-60 min., 4°C). The samples were neutralized and washed with FACS buffer (100 µL) twice prior to incubation in fixation buffer (100 µL, 15 min., 4°C). FACS buffer (100 µL) was added to neutralize the fixation buffer, the cells were centrifuged and resuspended in FACS buffer (200 µL). Flow cytometry was run using a BD FACSymphony A3 (BD Biosciences) and was analyzed using FlowJo.

### Ye infection

Ye Ruokola/71 (O:9) (13) was kindly provided by Dr. Mikael Skurnik at the University of Helsinki and used in this study. The strain was sequenced by SeqCenter (Pittsburgh, PA), annotated, and deposited in the NCBI database (Accession ID: JBXYBB000000000). To distinguish Ye from intestinal commensal bacteria, we generated a kanamycin resistant Ye strain by electroporating the low-copy plasmid carrying a kanamycin resistance gene (pYA4454-Kan^r^), into Ye Ruokola/71 (13), designated Ye Ruokola/71 K^+^. Electroporated cells were then incubated in Super Optimal broth with Catabolite repression (SOC media) (500 µL, 5 hrs., 25°C), spread on 25 µg/mL kanamycin Luria Broth (LB) agar plates, and incubated at 28°C for two days. A Ye Ruokola/71 K^+^ colony was cultured and stored in a -80 °C freezer.

One day prior to infection, Ye Ruokola/71 K^+^ (one loop) was inoculated from glycerol stock in 50 µg/mL kanamycin LB for overnight culture (3 mL, 28°C). The day of infection, the optical density (O.D.) of the overnight culture was measured, adjusted to 0.1 in 50 µg/mL kanamycin LB, and grown (15 mL, 28°C) until the O.D. reached 0.8. Then, the Ye culture was centrifuged (4,000 rpm, 10 min, 4°C), resuspended in sterile 1×PBS (1×, 3 mL), and diluted to the desired infection dosage.

Before infection, mice were deprived of food and water for six hours. Then, animals were administered Ye Ruokola/71 K^+^ (200 µL, 1×10^9^ colony forming units (cfu) by oral gavage (o.g.). The weight change and mortality of the infected mice were monitored for 21 days post infection (dpi). Because some mice that have marked weight loss are able to gain the weight back and recover from infection, we do not censor infected mice based off of their weight loss and use death as an endpoint for our experiments post Ye infection.

### Bacterial burden

Spleen, liver, and small intestine (∼50 mg) were collected from infected B6, 2wk Fe, and 4wk Fe mice. The tissue samples were homogenized in sterile 1×PBS (900 µL), diluted, and plated on 25 µg/mL kanamycin LB plates for each group of mice. The plates were incubated at 28°C for two days and the cfu of Ye Ruokola/71 K^+^ in each organ was compared accordingly.

### Iron chelation experiments

4wk Fe mice were administered 4 mg Deferoxamine (DFO) intraperitoneally (i.p.) or 10 mg/kg body weight Deferisarox (DFX) by o.g. every day for seven days. On the seventh day, serum was collected to verify whether the serum iron concentration decreased in the treated mice. On the eighth day, mice were infected with Ye Ruokola/71 K^+^ as described previously.

### Statistical analysis

Each experiment included a significant number of biological replicates to establish reproducibility. Statistical analyses of comparisons of data among groups were performed with one-way ANOVA/univariate or two-way ANOVA with Tukey post hoc tests. The log-rank (Mantel-Cox) test was used for survival analysis. All data were analyzed using GraphPad PRISM 10.1.2 software. The data are represented as the mean ± standard deviation (ns, no significance; * *P*< 0.05; ** *P*< 0.01; *** *P*< 0.001; **** *P*<0.0001).

## Results

### Exposure to high iron diet causes hemosiderosis in mice

To generate a dietary IO mouse model, C57BL/6 mice were fed a 2% carbonyl iron 10% kcal fat diet (high iron diet, HID) for two weeks (2wk Fe) or four weeks (4wk Fe) (Fig. 1A). Control groups were age and sex matched C57BL/6 (B6) mice, which remained on the Prolab IsoPro RMH 3000 5P76 diet (5P76 diet) provided by our animal resource facility. Overall, these two diets have similar ingredients; however, the 5P76 diet is a grain based, natural ingredient diet, while the HID uses highly refined ingredients. Table S1 compares the complete composition of the two diets and highlights the difference in iron concentration, in which the 5P76 diet contains 360 ppm and the HID contains 20037.1 ppm iron.

**Figure 1.**
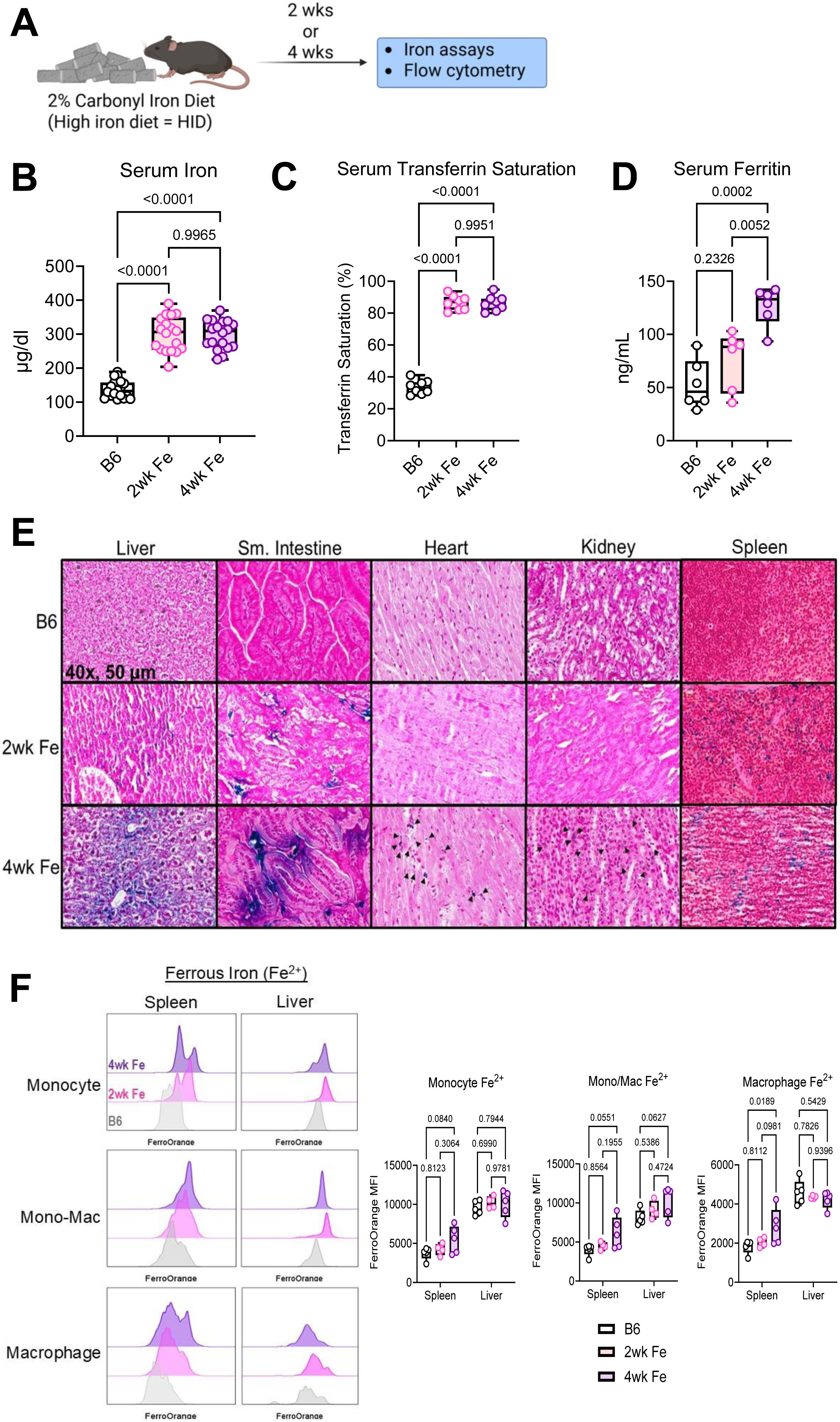
Exposure to high iron diet causes hemochromatosis in mice. (A) Schematic for generating iron overloaded mice using a 2% carbonyl iron diet. (B) Comparison of the serum iron content in WT B6 mice (B6), and mice on the high iron diet for 2 weeks (2wk Fe) and 4 weeks (4wk Fe) (*n* = 18 mice/group). Statistical analysis by ordinary one-way ANOVA with multiple comparisons. (C) Comparison of the transferrin saturation in B6, 2wk Fe, and 4wk Fe mice (*n* = 8 mice/group). Statistical analysis by ordinary one-way ANOVA with multiple comparisons. (D) Comparison of the serum ferritin concentration in B6, 2wk Fe, and 4wk Fe mice (*n* = 6 mice/group). Statistical analysis by ordinary one-way ANOVA with multiple comparisons. (E) Prussian blue staining for iron was performed on liver, small intestine, heart, and kidney sections from B6, 2wk Fe, and 4wk Fe mice. Images are shown at 40x, scale: 50µm. (ns, no significance; \**P* < 0.05; \*\**P* < 0.01; \*\*\**P* < 0.001; \*\*\*\**P* < 0.0001).

After two weeks and four weeks on the HID, mice had significantly increased serum iron concentrations compared to B6 mice (Fig. 1B). However, the serum iron concentration was similar for the 2wk Fe and 4wk Fe mice (Fig. 1B). Transferrin has two high-affinity binding sites for ferric iron, Fe^3+^, and is responsible for trafficking iron all over the body (29, 30). In healthy conditions, transferrin is 20-40% saturated, while transferrin saturation greater than 45% is an indicator of IO (31, 32). In agreement with this, the average transferrin saturation percentage was approximately 34% for B6 mice, while transferrin was about 86% saturated for 2wk and 4wk Fe mice (Fig. 1C). This indicates healthy iron metabolism in B6 mice and IO in the 2wk and 4wk Fe mice. Ferritin is an intracellular iron storage protein, which increases as time on the HID increases and is significantly higher in the 4wk Fe mice compared to B6 and 2wk Fe mice (Fig. 1D). These results suggest that as time on the HID increases, serum iron levels rise, causing excess iron to be stored intracellularly in ferritin nanocages.

High iron levels lead to the deposition of excess iron in various tissues (3). Staining for iron deposition showed that as time on the HID increases, there is increased tissue iron deposition (Fig. 1E). B6 mice have little to no visible iron deposited in the liver, small intestine, heart, kidney, or spleen. After two weeks on HID, iron begins to deposit in the small intestine, liver, and spleen, the sites of iron absorption, storage, and recycling, respectively. However, after four weeks on the HID, there is significant iron deposition in the liver, small intestine, and spleen, as well as detectable iron deposition in the heart and kidney (Fig. 1E).

During hemochromatosis hepcidin production is impaired, leading to decreased ferroportin degradation and continuous iron export from cells. In hemosiderosis, hepcidin levels increase, causing ferroportin to be degraded, and iron maintained intracellularly. To investigate this in our model, cells from the spleen and liver of B6, 2wk Fe, and 4wk Fe mice were isolated, stained for extracellular markers, and the level of intracellular ferrous iron, (Fe^2+^), was measured using FerroOrange. In general, there were higher levels of Fe^2+^ detected in MNPs of the liver than in the spleen for all three groups of mice (Fig. 1F). Levels of Fe^2+^ increase in MNPs as time on the HID increases in both the spleen and the liver (Fig. 1F). This data is in agreement with what would be expected for hemosiderosis rather than hemochromatosis. Altogether, the results suggest that keeping C57BL/6 mice on a HID is able to recapitulate the iron accumulation that is described in human hemosiderosis (3, 31).

### Systemic impacts of iron overload

Immune cell function and iron levels are closely linked. Therefore, we next characterized impacts of HID on the immune system. To investigate the systemic impacts of the HID, whole blood and serum were collected from B6, 2wk Fe, and 4wk Fe mice for a complete blood count (CBC) and to measure serum cytokines (Fig. 2A). The total number of white blood cells (WBCs) was significantly increased in the 4wk Fe mice compared to the other two groups, however there was no significant difference between the B6 mice and the 2wk Fe mice (Fig. 2B). Similar to the trend of total WBCs, the number of lymphocytes significantly increased in 4wk Fe mice compared to 2wk Fe and B6 mice, with no difference between the latter groups (Fig. 2B). For myeloid cells, the monocyte and neutrophil count increased as time on the HID increased (Fig. 2B&C), the number of eosinophils was significantly increased at four weeks on the HID, and there was no change in the number of basophils between the three groups (Fig. 2C).

**Figure 2.**
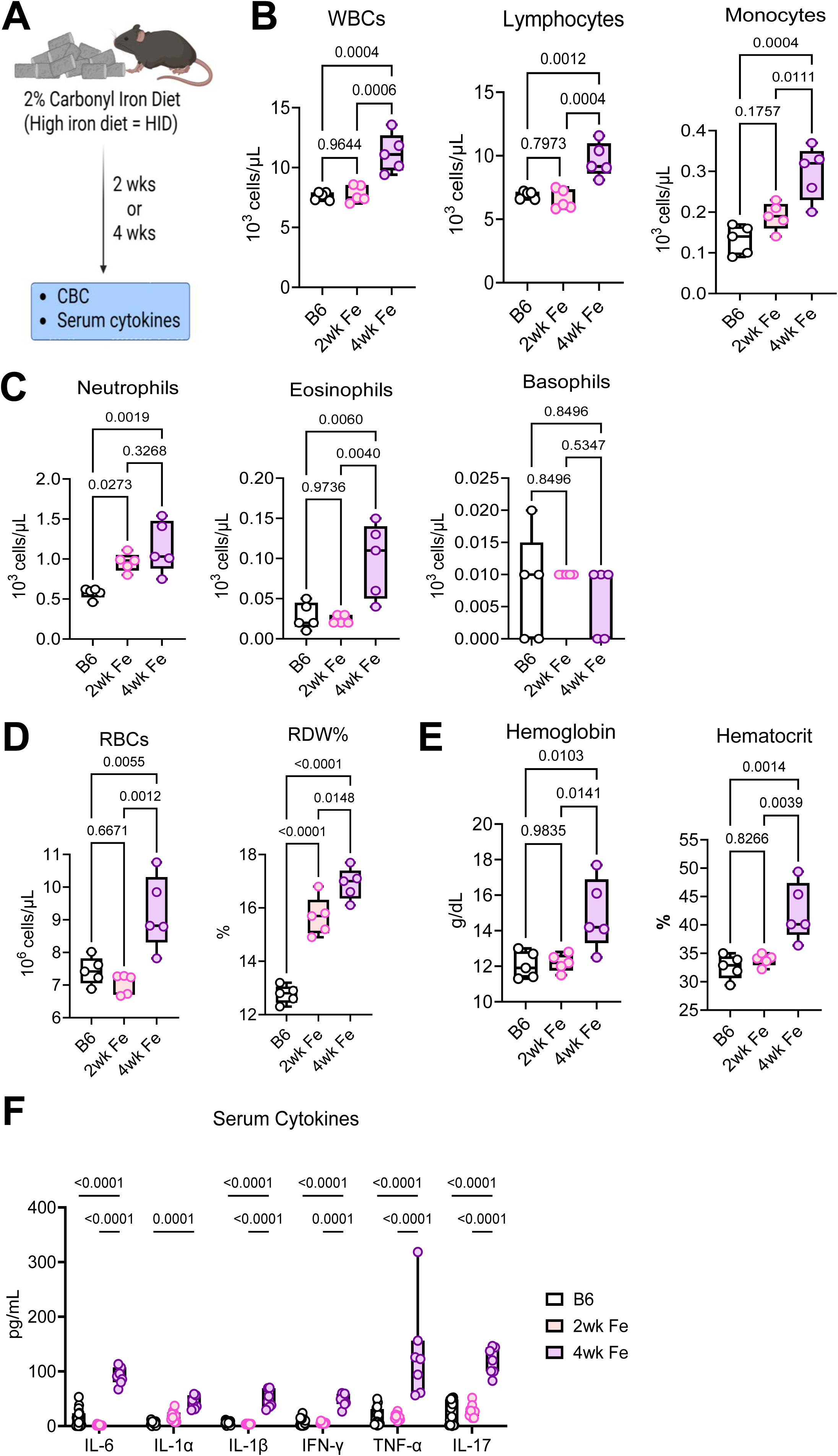
Impact of iron overload on the immune system. Complete blood count (CBC) analysis comparing the (A) the number of white blood cells (WBCs), lymphocytes, monocytes, and granulocytes, (B) the number of red blood cells (RBCs) and the red cell distribution width (RDW%), and (C) the hemoglobin and hematocrit levels between B6, 2wk Fe, and 4wk Fe mice (*n* = 5 mice/group). Statistical analysis by ordinary one-way ANOVA with multiple comparisons. (D) Comparison of serum cytokines in B6, 2wk and 4wk Fe mice (*n* = 17 B6, 10 2wk Fe, and 7 4wk Fe mice/group). Statistical analysis by two-way ANOVA with multiple comparisons. (ns, no significance; \**P* < 0.05; \*\**P* < 0.01; \*\*\**P* < 0.001; \*\*\*\**P* < 0.0001).

Other CBC parameters illustrated that after four weeks on the HID, mice had a significant increase in the number of red blood cells (RBCs) and the red cell distribution width (RDW%) increased as time on the HID increased (Fig. 2D). RDW% is a measurement of how varied the RBCs are in size and volume, and high RDW% can be indicative of various chronic diseases (33). Further, 4wk Fe mice have elevated hemoglobin and hematocrit compared to 2wk Fe and B6 mice (Fig. 2E), a phenomenon previously documented in individuals with IO (34, 35). Moreover, 4wk Fe mice have significantly increased concentrations of various inflammatory cytokines in the serum including interleukin-6 (IL-6), interleukin-1 (IL-1) family cytokines, interferon gamma (IFN-γ), tumor necrosis factor alpha (TNF-α), and interleukin-17 (IL-17) compared to 2wk Fe and B6 mice (Fig. 2F). Cytokines IL-1α and IL-17 follow the general trend that as time on the HID increases, serum IL-1α and IL-17 increase (Fig. 2F). For IL-6, IL-1β, IFN-γ, and TNF-α, there is no significant difference in serum cytokine level between the B6 and 2wk Fe mice, however there is a drastic increase in the serum cytokine level in the 4wk Fe mice (Fig. 2F).

### Impacts of iron overload on the intestine

The duodenum of the small intestine is the site responsible for iron absorption from the diet (36), and Ye is a food-borne pathogen that infects the intestine (11). Therefore, small intestine samples of B6, 2wk Fe, and 4wk Fe mice were collected to investigate the impacts of dietary IO on the intestine (Fig. 3A). It has been previously reported that iron deposition in the intestinal mucosa can cause physical changes to the intestinal epithelial barrier (IEB), activate immune responses within the lamina propria, and disrupt intestinal homeostasis (37). To determine whether the HID impacts the architecture of the small intestine, sections of small intestine samples from B6, 2wk Fe, and 4wk Fe mice were collected and stained with hematoxylin and eosin (H&E). Compared with B6 controls, 2wk Fe mice had mild inflammatory cell infiltration in the mucosal layer of the small intestine and minimal hyperplasia (Fig. 3B & Table 1). 4wk Fe mice had moderate inflammatory cell infiltration into both mucosal and submucosal layers of the intestine (Fig. 3B & Table 1). Further, 4wk Fe mice had moderate hyperplasia, marked goblet cell loss, and mild cryptitis (Fig. 3B & Table 1). According to the histopathological scoring, the HID causes progressive, time-dependent inflammatory cell infiltration and epithelial changes in the small intestine of mice, with little to no changes in mucosal architecture (Fig. 3B & Table 1).

**Figure 3.**
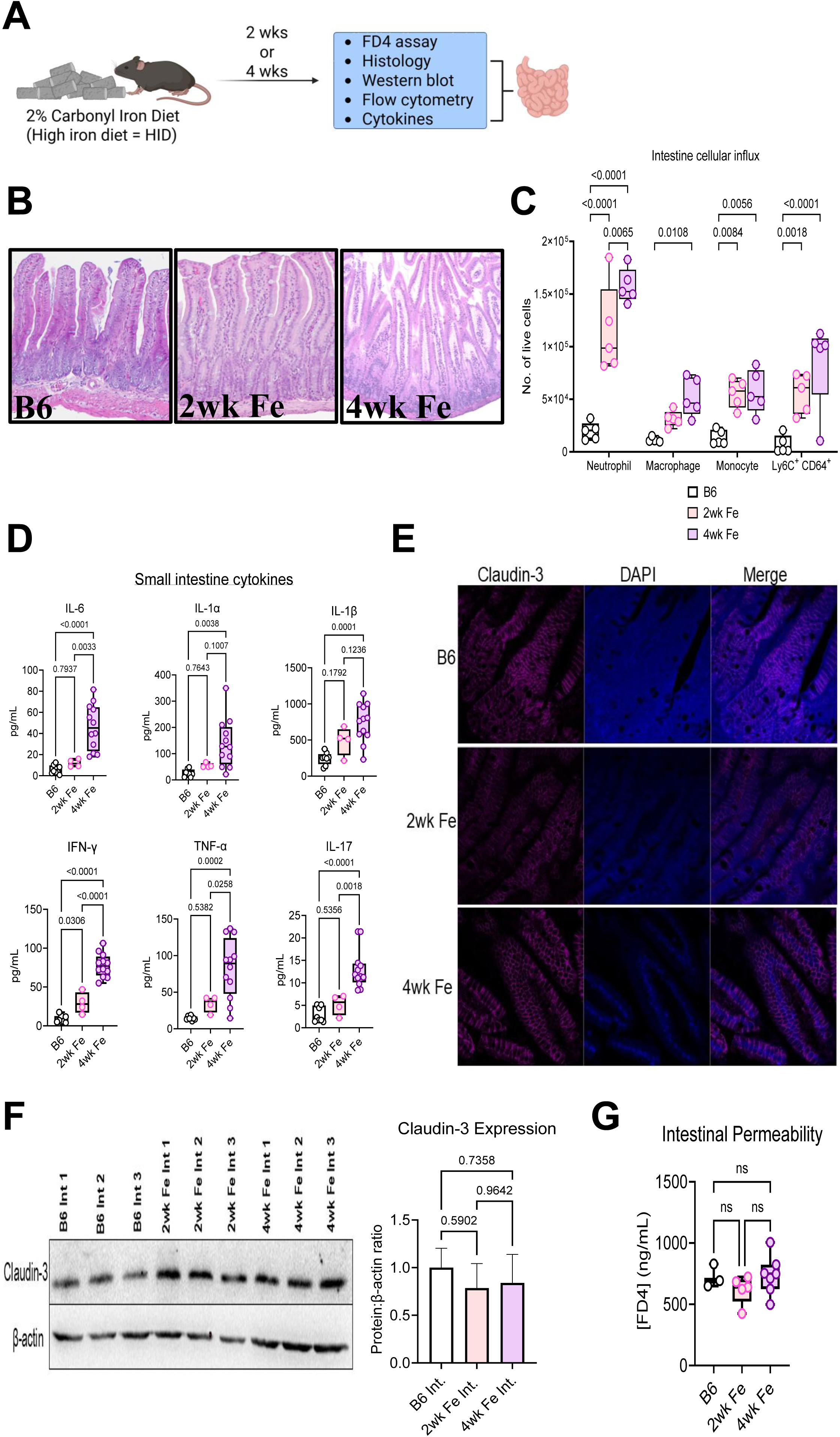
Impact of iron overload on the intestine. (A) Histopathology images (H&E stain) of small intestine sections of B6, 2wk Fe, and 4wk Fe mice. Images shown at 10x, scale: 250 µm. (B) Immunohistochemistry (IHC) of claudin-3 from B6, 2wk Fe, and 4wk Fe small intestine sections. Images shown at 40x, scale: 20 µm. (C) Comparison of the cellular influx in the intestine between B6, 2wk Fe, and 4wk Fe mice determined by flow cytometry. Gating strategy is as follows: neutrophils (CD45^+^ CD11b^+^ Ly6G^+^), macrophages (CD45^+^ CD11b^+^ Ly6G^-^ CD64^+^ Ly6C^-^), monocytes (CD45^+^ CD11b^+^ Ly6G^-^ CD64^-^ Ly6C^+^), and intermediate macrophages (CD45^+^ CD11b^+^ Ly6G^-^ CD64^+^ Ly6C^+^) (*n* = 5 mice/group). Statistical analysis by 2-way ANOVA with multiple comparisons. (D) Comparison of intestine cytokines in B6, 2wk, and 4wk Fe mice (*n* = 8 B6, 4 2wk Fe, and 12 4wk Fe). Statistical analysis by ordinary one-way ANOVA with multiple comparisons. (ns, no significance; \**P* < 0.05; \*\**P* < 0.01; \*\*\**P* < 0.001; \*\*\*\**P* < 0.0001).

**Table 1.**
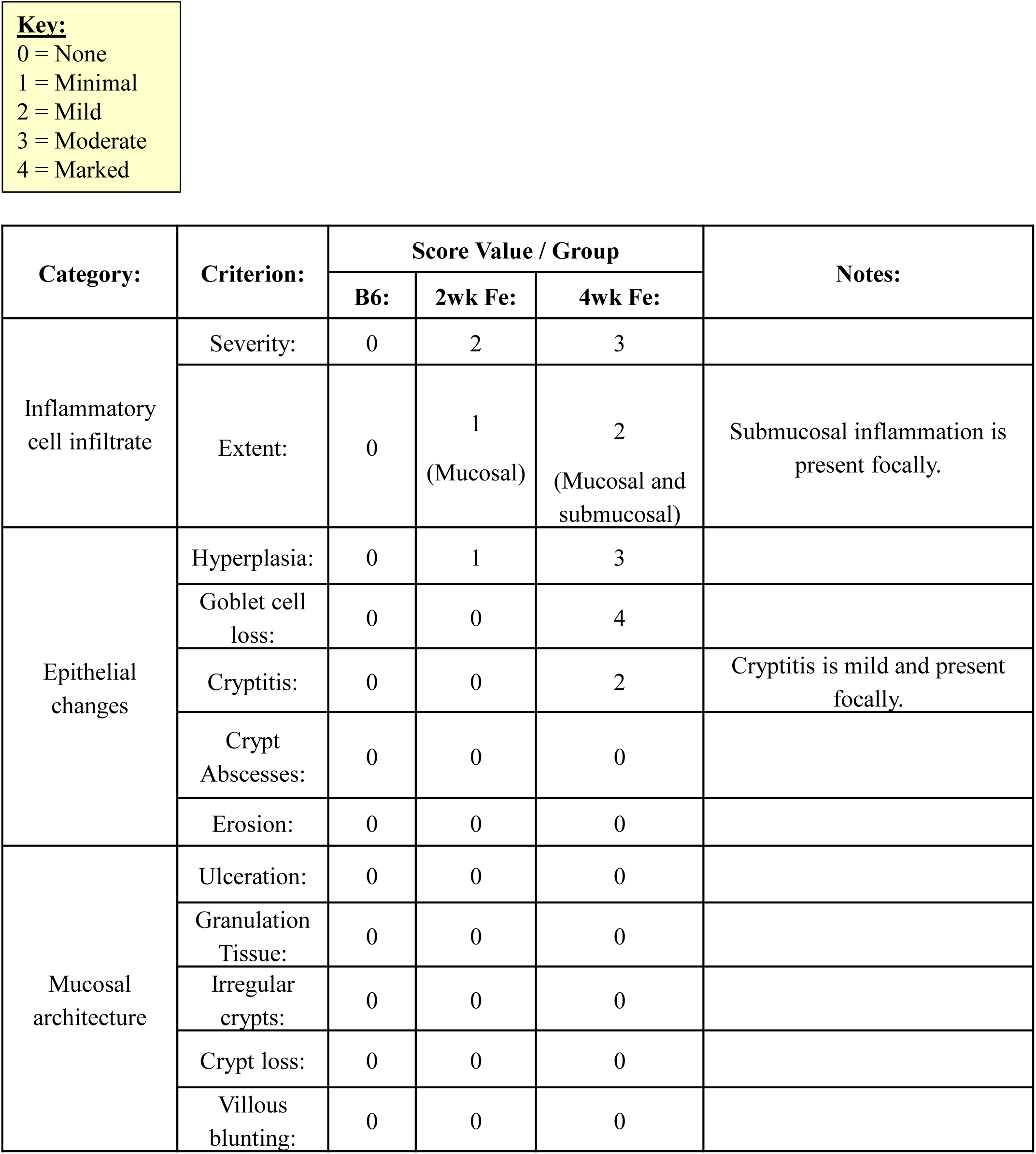

In agreement with the H&E staining, flow cytometry showed that IO causes increased immune cell recruitment to the intestine (Fig. 3C). 2wk Fe and 4wk Fe mice have significantly increased numbers of neutrophils, monocytes, and Ly6C^+^CD64^+^ MNPs in the small intestine compared to B6 mice (Fig. 3C). There is also an upward trend for the number of macrophages in the intestine of 2wk and 4wk Fe mice compared to the B6 mice (Fig. 3C). As observed in the serum, intestinal iron accumulation is associated with heightened levels of inflammatory cytokines (Fig. 3D). As time on the HID increases, the concentration of various inflammatory cytokines (IL-6, IL-1α, IL-1β, IFN-γ, TNF-α, and IL-17A) increases within the intestine, with 4wk Fe mice having the highest concentration of these cytokines compared to B6 and 2wk Fe mice (Fig. 3D).

It has also been reported that iron deposition in the intestinal mucosa can impair tight junction (TJ) proteins (37, 38). To test this in our model, we investigated whether there were differences in the expression of claudin-3, a component of TJ proteins, using immunohistochemistry (IHC). Results showed that claudin-3 was highly expressed and localized between intestinal epithelial cells along the villi in the small intestines of B6, 2wk Fe, and 4wk Fe mice (Fig. 3E). Interestingly, both IHC and western blot analyses indicated no significant differences in claudin-3 expression among the three groups (Fig. 3E and F). To further assess intestinal barrier function, a FITC-dextran assay was performed to measure intestinal permeability (Fig. 3G). No significant differences in intestinal permeability were observed among B6, 2wk Fe, and 4wk Fe mice. These findings are consistent with the claudin-3 data, as changes in claudin-3 expression would be expected to potentially reflect alterations in intestinal barrier integrity and permeability. Therefore, although 4wk-HID treatment induces changes in the IEB and triggers intestinal inflammation, it does not appear to significantly alter intestinal permeability at this point. Overall, HID causes IO that drives systemic inflammation characterized by increased WBCs, abnormalities in RBCs, and significantly elevated inflammatory cytokine concentrations in the serum of iron-loaded mice. Further, iron deposition in the intestine alters the IEB, leading to increased immune cell influx into the intestine and significantly increased levels of inflammatory cytokines. Therefore, the results suggest that IO causes systemic and intestinal inflammation.

### Hemochromatosis causes increased susceptibility to *Yersinia* infection

To study the impact of dietary IO during Ye infection, B6, 2wk Fe, and 4wk Fe mice were infected orally with Ye Ruokola/71 K^+^ (Fig. 4A). Mice were deprived of food and water for six hours prior to infection. This is done to ensure mice have an empty stomach for facilitating bacterial colonization and adherence to the intestinal epithelium. Mouse survival was monitored for twenty-one dpi (Fig. 4A). Ye Ruokola/71 is considered weakly pathogenic and is unable to kill mice unless immunocompromised, such as in the case of IO (11, 39). In agreement with this, survival studies showed that B6 mice completely survived infection with 1×10^9^ cfu Ye Ruokola/71 K^+^ (Fig. 4B). In contrast, 2wk and 4wk Fe mice succumbed to Ye Ruokola/71 K^+^ infection within two weeks (Fig. 4B). Upon Ye infection, all three groups of mice lose weight for the first four days, however by six dpi the B6 mice begin to regain their weight, indicating that the B6 mice are recovering at this time point (Fig. 4C). In the iron overloaded mice, their weight continues to decrease overtime until the mice succumb to infection (Fig. 4C).

**Figure 4.**
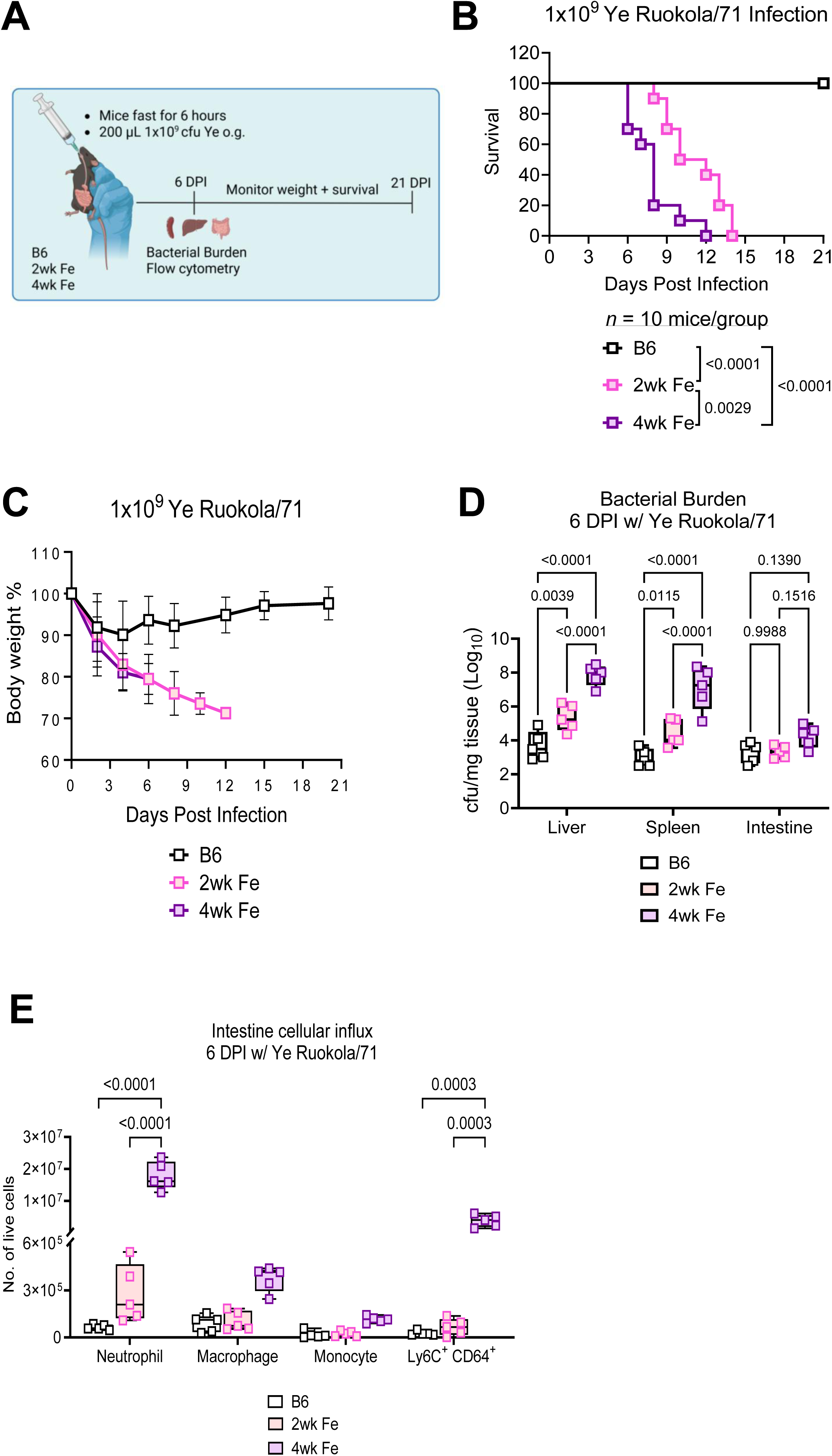
Hemochromatosis causes increased susceptibility to *Yersinia* infection. (A) Schematic for Ye-infection experiments. (B) Survival of B6, 2wk Fe, and 4wk Fe mice infected with 1x10^9^ cfu of Ye Ruokola/71 (*n* = 10 mice/group). Statistical analysis by the Log-rank (Mantel-Cox) test. (C) Weight change of B6, 2wk Fe, and 4wk Fe mice post Ye-infection (*n* = 5 B6, 10 2wk Fe, and 5 4wk Fe mice). (D) Comparison of the bacterial burden in the liver, spleen, and small intestine of B6, 2wk Fe, and 4wk Fe mice six days post infection with Ye Ruokola/71 K^+^ (*n* = 5 mice/group). Statistical analysis by 2-way ANOVA with multiple comparisons. (E) Comparison of the cellular influx in the small intestine between B6, 2wk Fe , and 4wk Fe mice six days post Ye Ruokola/71 infection determined by flow cytometry; gating strategy as described previously (*n* = 5 mice/group). Statistical analysis by 2-way ANOVA with multiple comparisons. (ns, no significance; \**P* < 0.05; \*\**P* < 0.01; \*\*\**P* < 0.001; \*\*\*\**P* < 0.0001).

At six dpi, the B6 mice are recovering, while the iron-overloaded mice continue to worsen. Therefore, we chose this timepoint to evaluate differences in bacterial burden and immune cell infiltration. At six dpi, the 2wk Fe and 4wk Fe mice have significantly higher bacterial burden detected in the liver and spleen, however there is no difference in the small intestine (Fig. 4D). This result may be because at this timepoint, the Ye has disseminated out of the small intestine via the Peyer’s Patches to the spleen and liver. Additionally, the 4wk Fe mice exhibited the highest bacterial burden among the groups, significantly greater than 2wk Fe mice in the liver and spleen (Fig. 4D). Although bacterial levels in the small intestine were similar, 4wk Fe mice showed marked intestinal inflammation at 6 dpi, with increased infiltration of neutrophils, Ly6C CD64 MNPs, macrophages, and monocytes compared to 2wk Fe and B6 mice (Fig. 4E). In 2wk Fe mice, neutrophil numbers were elevated relative to B6 mice, but mononuclear phagocyte populations were comparable between these groups (Fig. 4E). The higher bacterial burden and intestine cellular influx in the 4wk Fe mice provides a reason for why this group succumbs to infection significantly faster than the other two groups of mice (Fig. 4B). These results suggest that IO hosts are susceptible to Ye infection, which is attenuated in B6 mice.

### Impact of iron chelation on Ye infection

Deferoxamine (DFO) and Deferisarox (DFX) are treatment options for removing excess iron in individuals with IO disorders (3). These iron chelators bind to excess iron in tissues and form complexes that are excreted from the body. Therefore, we tested the impact of iron chelation therapy on 4wk Fe mice during Ye infection because this group experienced the most iron deposition in tissues (Fig. 1E). 4wk Fe mice were administered DFO, DFX, or PBS as a control for seven days prior to infection with Ye (Fig. 5A). For the seven days of treatment and infection, the HID was removed and replaced with the 5P76 chow diet provided by the animal resource facility (Fig. 5A). The diet was removed because if the mice remained on the HID while receiving iron chelation therapy, their serum iron levels did not decrease (data not shown). To test whether the serum iron levels decreased post iron chelation treatment, serum was collected prior to and post treatment (Fig. 5A). After treatment with DFO and DFX, the serum iron concentration was significantly less than the concentration prior to treatment (Fig. 5B). However, the serum iron concentration in the 4wk Fe + PBS group also decreased post treatment, indicating that removing the HID was enough to decrease the serum iron levels in iron overloaded mice (Fig. 5B). In fact, switching the diets dramatically altered the serum iron level and decreased the concentration to approximately 100 µg/dl, less than the average serum iron concentration in B6 mice (Fig. 1B). Mice and rats have a significant capacity for iron excretion, differing from humans that have no significant physiological mechanisms for eliminating excess iron, causing it to accumulate in the body (40, 41). Upon infection with Ye, the 4wk Fe + PBS group had 100% survival (Fig. 5C), with only minimal weight loss during the first few days of infection and made a full recovery by 21 dpi (Fig. 5D). In contrast, 4wk Fe mice treated with DFO had severe weight loss post infection and succumbed to infection within 9 days (Fig. 5C & D), which is sooner than both the 2wk Fe and 4wk Fe groups (Fig. 4B). DFX treated FeB6 mice also had increased mortality compared to the PBS treated group (Fig. 5C), indicating that iron chelation treatment is detrimental during Ye infection. These results indicate that removal of the HID for a week decreases the serum iron concentration in 4wk Fe mice and rescues them from Ye infection. However, treatment of these mice with DFO or DFX is detrimental during Ye infection, evidenced by increased mortality in these groups.

**Figure 5.**
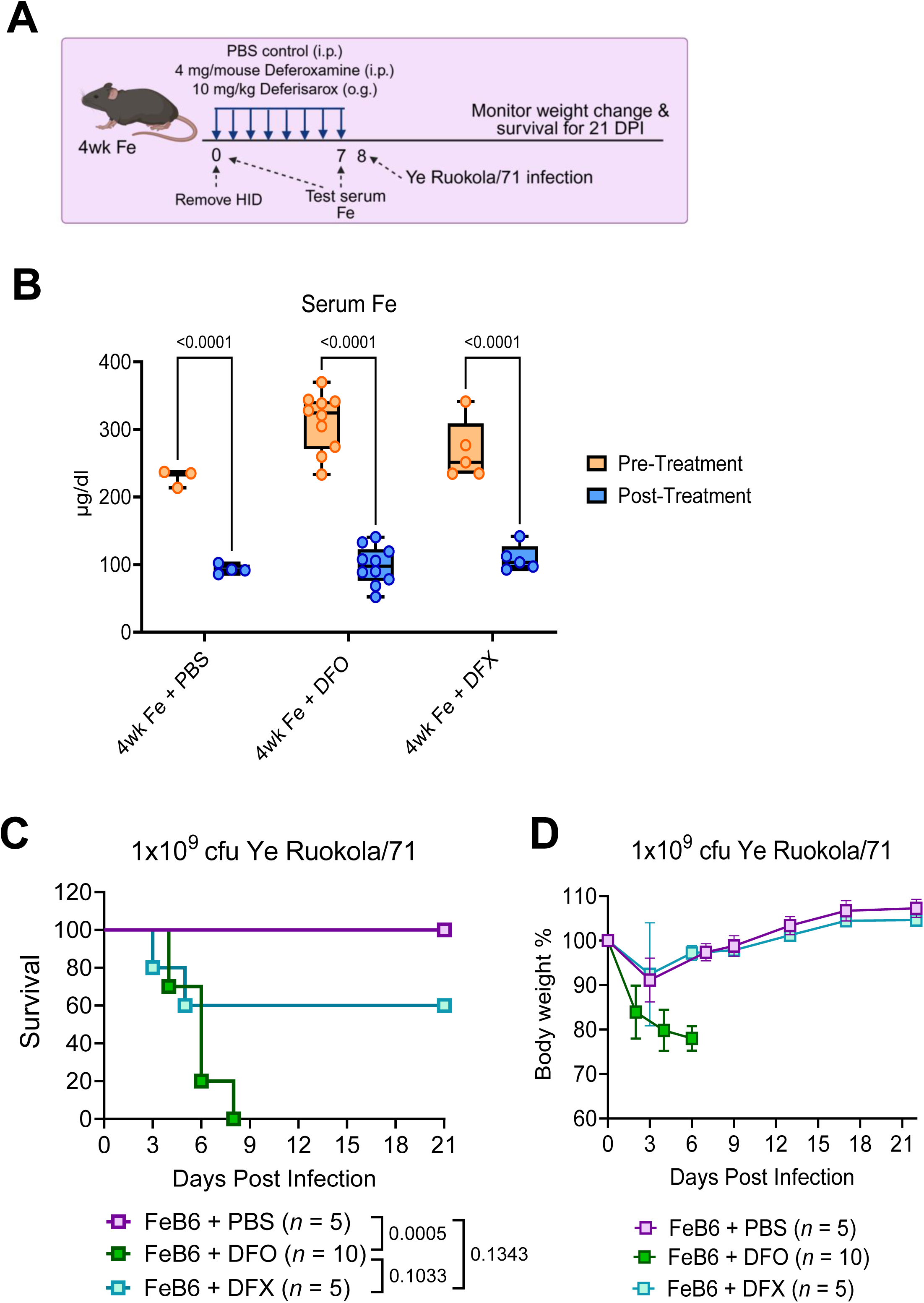
Impact of iron chelation during Ye infection. (A) Schematic of iron chelation treatment by Deferoxamine (DFO), Deferisarox (DFX), or phosphate buffered saline (PBS) as a control group. (B) Comparison of the serum iron levels in 4wk Fe (FeB6) mice pre- and post-treatment with PBS, DFO, or DFX (*n* = 3 PBS pre-treatment and 4 PBS post-treatment, 10 DFO, and 5 DFX-treated mice). Statistical analysis by two-way ANOVA with multiple comparisons. (C) Concentration of hepcidin in the serum of B6 and FeB6 mice (*n* = 4 mice/group). Statistical analysis by unpaired t-test. (D) Survival of FeB6 mice treated with PBS, DFO, or DFX infected with 1x10^9^ cfu of Ye Ruokola/71 (*n* = 5 PBS, 10 DFO, and 5 DFX-treated FeB6 mice). Statistical analysis by Log-rank (Mantel-Cox) test. (E) Weight change of PBS, DFO, and DFX-treated FeB6 mice post Ye-infection (*n* = 5 PBS, 10 DFO, and 5 DFX-treated FeB6 mice). (ns, no significance; \**P* < 0.05; \*\**P* < 0.01; \*\*\**P* < 0.001; \*\*\*\**P* < 0.0001).

## Discussion

Here, we characterize a dietary IO mouse model that recapitulates the iron accumulation, immune dysregulation, and susceptibility to Ye infection that is observed in humans with hemosiderosis. Although hemochromatosis and hemosiderosis have been shown to cause similar outcomes during bacterial infections, the underlying mechanisms may differ because, unlike hemochromatosis, hemosiderosis is typically characterized by increased hepcidin levels (42).

In type 1 hemochromatosis, caused by mutations in the *Hfe* gene encoding the human homeostatic iron regulator protein, iron accumulation is chronic. Humans are not capable of removing excess iron, causing IO, and eventually leading to worsening organ damage and associated pathologies over time. Since females lose iron during menstruation, they become symptomatic later in life than males and develop disease in their sixties (3, 43). Conversely, males are affected 2-3 times more than females and develop the disease in their fifties (3, 43). *Hfe^tm2Nca^* mice, which are homozygous for a targeted mutation of *Hfe*, are used to model type I hemochromatosis (38, 44). When using the *Hfe^tm2Nca^* mouse model, it is important to consider that this model is similar to human hemochromatosis in that IO is chronic, so the mice do not experience symptoms until later in life. In general, an “older mouse,” about 18-24 months old, correlates to a human in the age range of 56-69 years old (45). Therefore, IO studies using *Hfe^tm2Nca^* mice might be informative when the mice are about 18-24 months old. However, this still may not be old enough to recapitulate the metabolic disruption that occurs from iron accumulation in human tissues over decades (46). Additionally, mice at this age begin to experience senescence and the onset of strain-specific diseases that can be misleading when interpreting data (45). Further, aging can greatly impact immune cell number, function, and outcomes post infection in mice (43). Lastly, it should be considered that sex differences impact hepcidin expression, iron storage, and the severity of diseases that result from IO in both mice and humans (47). In our dietary IO model, it appears that sex differences are minimized, as similar numbers of male and female mice were used in this study, and we do not find much variation among these data points. This could be due to the fact that female C57BL/6 mice have an estrus cycle rather than a menstrual cycle (48), which may minimize the difference in iron accumulation among sexes that is observed in humans. However, it would be important to confirm whether there are sex differences in *Hfe^tm2Nca^*mice when being used in studies. Since hemosiderosis and hemochromatosis are different, results outlined here need to be corroborated in hemochromatosis mouse models to determine how impacts of IO on immunity and Ye infection are similar in these IO disorders and how they are different.

Disorders of iron metabolism are often linked to intestinal tissue damage, leading to dysfunctional homeostasis and immunity in the intestine (37). It has been reported that IO causes mucosal injury leading to cell death (37), destruction of epithelial TJs (49), lipid peroxidation, and intestinal inflammation (50). Further, IO promotes enhanced neutrophil recruitment (38, 51), impairs monocyte-macrophage differentiation (52), and polarizes macrophages to an inflammatory M1 phenotype while inhibiting M2 macrophage polarization (53). This can exacerbate the intestinal inflammation induced by IO. In agreement with our study, other studies have seen that IO upregulates pro-inflammatory cytokines, including IL-1β and IL-6, while downregulating anti-inflammatory cytokines, including IL-4 and IL-10, in the intestine of mice (49). During homeostasis, the intestine maintains a tolerogenic environment through the production of anti-inflammatory cytokines produced by macrophages and regulatory T cells (Tregs) (54, 55). Since IO polarizes macrophages to be more inflammatory and causes Treg cell death (51), perhaps this contributes to the excessive intestinal inflammation observed during IO. Thus, further investigation is needed to characterize the immune cell dysfunction identified in the intestine.

Ye is a heterogeneous group classified into six biovars and numerous serotypes. Serotypes O:3 (biovar 4), O:5 (biovars 2 and 3), O:8 (biovar 1B), and O:9 (biovar 2) are most frequently isolated from human samples worldwide (56). High-virulence O:8 strains are more common in the United States, whereas the lower-virulence O:3 and O:9 strains predominate in Europe (57). The major distinction between O:8 and O:9 strains is the presence or absence of the high-pathogenicity island (HPI), a chromosomal region encoding the siderophore yersiniabactin (11, 39, 58), which facilitates Fe³ acquisition from host proteins (58). Intriguingly, the O:8 strain WA (ATCC 27729) caused similar mortality in WT and iron-overloaded Hepcidin knockout (HKO) mice (14). In contrast, the O:9 strain Ruokola/71 was more lethal in dietary IO mice (Fig. 4B) and HKO mice (14), than in WT B6 mice. Weakly pathogenic Ruokola/71 (O:9) causes complete mortality in dietary IO mice, suggesting the severe impairment of HID on host immunity rather than the iron-scavenging capabilities and virulence of the bacteria. We reason that differences in lethality may be due to bacterial strain-specific differences, infection dose, or iron accumulation between WT B6, dietary IO, HKO, and *Hfe^tm2Nca^* mice and plan to assess differences between O:9 and O:8 infections in future studies. In addition, individuals with IO are susceptible to various kinds of enteric bacteria aside from *Yersinia*, including *Salmonella typhimurium* (26, 59), *Klebsiella pneumoniae* (60), and *Vibrio vulnificus* (61, 62), and these species often produce siderophores as well (63–65). In future studies, it is important to evaluate the impact of dietary IO on siderophore-producing Ye strains as well as other enteric pathogens.

There is no cure for IO disorders, however it can be treated by therapeutic phlebotomy or iron chelation treatments (31). In our iron chelation experiments, the HID had to be replaced with the animal facility’s chow diet during treatment. This was necessary because if the mice remained on the HID during treatment, their serum iron levels did not decrease (data not shown). Interestingly, we observed that replacing the diet for seven days was enough to reverse the significantly high serum iron concentration observed in the 4wk Fe mice. This result was surprising because it has been reported that mammals lack strong mechanisms for iron excretion, and rather iron homeostasis is largely dictated by iron absorption from the diet (40). Reasons for this could be due to differences in iron metabolism between mice and humans (40, 41). Previous studies have indicated that unlike humans, mice have mechanisms for removing excess body iron via the gastrointestinal tract and urinary system, and iron overloaded mice have increased rates of iron excretion (40, 41). This could indicate why the serum iron level in the mice drastically decreased after the iron source was removed.

We also observed that iron chelation drugs DFO and DFX are detrimental during Ye infection. While it has been reported that DFO treatment promotes Ye infection in humans due to enhancements in growth and virulence (66), there is a lack of data regarding the impacts of DFX on Ye infection. Further investigation is required to determine the impacts of iron chelators on host immunity and Ye infection, and this is an important area of research to pursue since individuals with IO are more susceptible to Ye infection and also may be receiving these treatments.

Altogether, this study demonstrates that maintaining mice on a HID for two or four weeks could recapitulate the iron accumulation, immune dysfunction, and infection susceptibility observed in humans with hemosiderosis. This dietary IO mouse model can be utilized to characterize the impacts of hemosiderosis on immunity and how these impacts exacerbate enteric infection. Such advances will aid in improving the prevention and treatment of enteric infections in individuals with IO.

## Acknowledgments

We thank Dr. Mikael Skurnik from the University of Helsinki for kindly providing *Y. enterocolitica* Ruokola/71. This work was partially supported by the National Institutes of Health grant R01AI162670 to WS. We’d like to thank the Animal Resource Facility and the immunology core within the Department of Immunology and Microbial Disease at Albany Medical College.

## Contributions

Experiments conceived and designed by WS, MV, and SD; experiments performed by MV and SD; data analyzed by MV, SD, NV, DZ, and WS; manuscript written by MV; and manuscript edited by WS.

