## Supplementary material for "Dietary iron overload enhances susceptibility to *Yersinia enterocolitica* infection": Supplemenatary

### Table S1

**Key:**
B6 diet: Prolab IsoPro RMH 3000 5P76
Fe diet: 2% Carbonyl Iron 10% kcal Fat Diet (93G)

**Macronutrients.**

| Calories from: | B6 Diet | Fe Diet |
|---|---|---|
| Protein: | 26.126% | 19.700% |
| Carbohydrates: | 59.497% | 70.300% |
| Fat: | 14.377% | 10.000% |

**Minerals.**

| | B6 Diet | Fe Diet |
|---|---|---|
| Calcium | 1.09% | 0.50% |
| Phosphorus | 0.79% | 0.30% |
| Phosphorus (non-phytate) | 0.47% | X |
| Potassium | 0.95% | 0.36% |
| Sodium | 0.22% | 0.10% |
| Chlorine | 0.40% | 0.16% |
| Magnesium | 0.23% | 0.0519% |
| Sulfur | 0.29% | X |
| Copper | 14 ppm | 6.2 ppm |
| Iron | 360 ppm | 20037.1 ppm |
| Zinc | 110 ppm | 41.6 ppm |
| Manganese | 100 ppm | 10.5 ppm |
| Iodine | 0.99 ppm | 0.21 ppm |
| Selenium | 0.41 ppm | 0.15 ppm |
| Molybdenum | Information not provided. | 0.15 ppm |
| Chromium | 0.01 ppm | 1.00 ppm |
| Fluorine | 17 ppm | X |
| Cobalt | 0.41 ppm | X |

**Amino Acids.**

| | B6 Diet | Fe Diet |
|---|---|---|
| Lysine | 1.31% | 1.40% |
| Methionine | 0.58% | 0.46% |
| Cystine | 0.40% | 0.35% |
| Arginine | 1.41% | 0.66% |
| Phenylalanine | 0.99% | 0.88% |
| Tyrosine | 0.64% | 0.92% |
| Histidine | 0.56% | 0.50% |
| Isoleucine | 0.90% | 1.0% |
| Leucine | 1.64% | 1.60% |
| Threonine | 0.82% | 0.76% |
| Tryptophan | 0.28% | 0.20% |
| Valine | 1.03% | 1.20% |
| Aspartic Acid | 2.30% | 1.20% |
| Glutamic Acid | 5.06% | 3.64% |
| Alanine | 1.20% | 0.52% |
| Glycine | 1.11% | 0.32% |
| Proline | 1.49% | 1.80% |
| Serine | 1.12% | 1.0% |
| Taurine | 0.03% | X |

**Vitamins.**

| | B6 Diet | Fe Diet |
|---|---|---|
| Vitamin A | 18000 IU/kg | 6000 IU/kg |
| Vitamin D | 2500 IU/kg | 1500 IU/kg |
| Vitamin E | 75 IU/kg | 113 IU/kg |
| Vitamin K | 1.9 ppm | 3.1 ppm |
| Biotin | 0.40 ppm | 0.30 ppm |
| Choline | 1730 ppm | 1262.3 ppm |
| Folic Acid | 1.2 ppm | 3.0 ppm |
| Niacin | 60 ppm | 45.0 ppm |
| Pantothenate/ Pantothenic Acid | 13 ppm | 22.0 ppm |
| Riboflavin | 14 ppm | 9.0 ppm |
| Thiamin | 9.7 ppm | 7.3 ppm |
| Vitamin $B_6$ / Pyridoxine | 8.3 ppm | 8.6 ppm |
| Vitamin $B_{12}$ | 0.077 ppm | 0.04 ppm |
| Vitamin C / Ascorbic Acid | 0.0 ppm | 0.0 ppm |
| Carotene | 1.2 ppm | X |

**Fatty Acids.**

| | B6 Diet | Fe Diet |
|---|---|---|
| 4:0 Butyric | X | 0.0 % |
| 6:0 Caproic | X | 0.0 % |
| 8:0 Caprylic | X | 0.0 % |
| 10:0 Capric | X | 0.0 % |
| 12:0 Lauric | X | 0.0 % |
| 14:0 Myristic | X | 0.0 % |
| 16:0 Palmitic | X | 0.42% |
| 16:1 Palmitoleic | X | 0.0% |
| 18:0 Stearic | X | 0.15% |
| 18:1 Oleic | X | 0.87% |
| 18:2 Linoleic | 1.60% | 2.0% |
| 18:3 Linolenic | 0.17% | 0.3% |
| Arachidonic Acid | 0.02% | X |
| Omega-3 Fatty Acids | 0.34% | X |

**Other.**

| | B6 Diet | Fe Diet |
|---|---|---|
| Cholesterol | 198.0 ppm | 40.0 ppm |
| Fiber | 4.3% | 3% |
